# Neuronal composition of processing modules in human V1: laminar density for neuronal and non-neuronal populations and a comparison with macaque

**DOI:** 10.1101/2023.05.21.540412

**Authors:** V Garcia-Marin, JG Kelly, MJ Hawken

## Abstract

The neuronal composition of homologous brain regions in different primates is important for understanding their processing capacities. Primary visual cortex (V1) has been widely studied in different members of the Catarrhines or Old-World monkeys. Neuronal density is considered to be central in defining the structure--function relationship. In human, there are large variations in the reported neuronal density from prior studies. We found the neuronal density in human V1 was 79,000 neurons/mm^3^, which is 35% of the neuronal density previously determined in macaque V1. Laminar density was proportionally similar between human and macaque. In V1, the ocular dominance column (ODC) contains the circuits for the emergence of orientation preference and spatial processing of a point image in many mammalian species. Analysis of the total neurons in an ODC and of the full number of neurons in macular vision (the central 15 degrees) indicate that humans have 1.28 times more neurons than macaques even though the density of neurons in macaque is 3 times the density in human V1. We propose that the number of neurons in a functional processing unit rather than the number of neurons under a mm^2^ of cortex is more appropriate for cortical comparisons across species.

## Introduction

It has been long established that the connectivity both within and between layers of cortex is an important feature enabling the understanding of the structure and function of cortical circuitry. Recent studies have emphasized the importance of determining the density of different neuron populations within cortical areas (Dombrowski et al, 2001; Charvet et al, 2015; von Bartheld et al, 2016) to understand uniformity principles within cortical areas and across cortical layers (Meyer et al, 2010; Wagstyl et al, 2018; Chariker et al, 2016, 2018, 2021, 2022) as a crucial step in determining the details necessary to make population models of cortical circuits.

Visual cortex is among the most extensively studied cortical areas across many mammalian species, including humans. The early visual pathway of different members of the Catarrhines or Old-World monkeys, the parvorder that includes macaque monkeys and the Great Apes, show considerable similarity (de Sousa et al, 2010; Kaas, 2020). The photoreceptors, post-receptoral neurons of the retina and those in main relay nucleus to cortex, the lateral geniculate nucleus (LGN), in the different subfamilies are closely matched (Boycott and Dowling, 1969; Kolb and DeKorver, 1991; Sumner and Mollen, 2000; de Sousa et al, 2013). The identification of thalamic terminal distributions using immunocytochemical labelling with the vesicular glutamate transporter (vGluT2) (Fremeau et al., 2001; Fujiyama et al., 2001) has made study of the afferents into human cortex and a comparison with macaque possible (Garcia-Marin et al, 2013). There are many similarities in vGluT2-ir distributions between macaque and human primary visual cortex, such as high densities of vGluT2-ir terminals in layer 4C, patches of VGluT2-ir puncta in the supragranular layers (2/3), lower densities but clear distributions in layers 1 and 6, and very few puncta in layers 5 and 4B (Garcia-Marin et al, 2013). Nonetheless, there are also some major differences. For example, the structure (Preuss et al, 2002) and presumptive thalamic input (Garcia-Marin et al, 2013) to layer 4A is quite distinct between macaque and human.

Studies of human V1 have reported substantially different neuronal densities, ranging from 18 x 10^3^ (Pakkenberg, 1966) to 123 x 10^3^ neurons/mm^3^ (Selemon et al, 1995; see suppl table 1 for additional studies). In general, studies that used non-stereological techniques to estimate neuronal density yielded overall densities that were lower than estimates of neuronal density using stereological methods (Everall et al, 1993; Pakkenberg and Gundersen, 1997; Inda et al, 2007; Dorf-Petersen et al, 2007; van Kann et al, 2017) or 3D counting methods (Selemon et al, 1995). However, even among the studies that used stereological or 3D counting methods the average densities varied considerably between studies, up to a factor of 3 (compare Selemon et al, 1995 and Inda et al, 2007, in suppl. Table 1). This variability in the results of the overall neuronal density of human V1 is also observed in the results from studies that focused on the laminar densities (see Supl Table 1). When laminar densities were determined for a single layer, large differences were observed in the estimates across studies; for example, in layer 4C estimates range from 47 to 195 x 10^3^ neurons/mm^3^ (a factor of 4: Inda et al, 2007; Van Kann et al, 2017; see suppl table 1 for other studies).

**Table 1:** Total number of cells (#DAPI) or neurons (#NeuN, #PV) for each of the 7 humans (H1-H7) in each sampling column (b1-b2). Volume of tissue analyzed (mm^3^), cortex length (mm) and number of optical planes taken for each human and column. Densities for DAPI, NeuN, and PV labelled cells, were calculated using the total number of cells (#DAPI, #Neun, #PV) and the volume (mm^3^) in each column. (A column of tissue was defined spanning from the pial surface to the white matter and oriented orthogonal to the pial surface)

| Human | #DAPI | Dens_Dapi<br>cell/mm<sup>3</sup> | #NeuN | Dens_NeuN<br>neurons/mm<sup>3</sup> | #PV | Dens_PV<br>PV/mm<sup>3</sup> | %PV | #<br>Planes | Vol<br>mm<sup>3</sup> | cx length<br>mm |
| --- | --- | --- | --- | --- | --- | --- | --- | --- | --- | --- |
| H1b1 | 1,049 | 145,855 | 480 | 66,740 | 25 | 3,476 | 5.2 | 12 | 0.007 | 2.77 |
| H1b2 | 1,915 | 142,812 | 814 | 60,704 | 38 | 2,834 | 4.7 | 22 | 0.013 | 2.82 |
| H2b1 | 2,388 | 156,827 | 1131 | 74,276 | 62 | 4,072 | 5.5 | 25 | 0.015 | 2.82 |
| H2b2 | 3,476 | 186,882 | 1825 | 96,053 | 93 | 5,000 | 5.1 | 29 | 0.019 | 3.00 |
| H3b1 | 2,903 | 181,125 | 1266 | 78,989 | 84 | 5,241 | 6.6 | 23 | 0.016 | 3.23 |
| H3b2 | 2,526 | 146,837 | 1216 | 70,686 | 68 | 3,953 | 5.6 | 27 | 0.017 | 2.95 |
| H4b1 | 1,749 | 166,949 | 859 | 81,995 | 29 | 2,768 | 3.4 | 18 | 0.010 | 2.69 |
| H4b2 | 1,035 | 168,858 | 498 | 81,248 | 34 | 5,547 | 6.8 | 12 | 0.006 | 2.36 |
| H5b1 | 853 | 152,727 | 419 | 75,021 | 27 | 4834 | 6.4 | 10 | 0.006 | 2.58 |
| H5b2 | 1,741 | 167,302 | 796 | 76,492 | 58 | 5,574 | 7.3 | 19 | 0.010 | 2.53 |
| H6b2 | 2,269 | 156,581 | 1204 | 83,087 | 103 | 7,108 | 8.6 | 30 | 0.014 | 2.23 |
| H6b3 | 1,071 | 197,919 | 534 | 98,682 | 29 | 5,359 | 5.4 | 9 | 0.005 | 2.78 |
| H7b2 | 2,450 | 173,348 | 1214 | 85,896 | 58 | 4,104 | 4.8 | 22 | 0.014 | 2.97 |
| **Mean** | **1,956** | **164,925** | **943** | **79,221** | **54** | **4,605** | **5.8** | **20** | **0.012** | **2.75** |
| SEM | 224 | 5,833 | 114 | 3,025 | 8 | 396 | 0.411 | 2.97 | 0.002 | 0.08 |

Although there have been comparisons between the neuronal density in humans and macaques, advances in neuron identification and large-scale imaging combined with automated image analysis have made it possible to make more rigorous quantitative comparisons. Recent studies using these methodological approaches (Kelly and Hawken, 2017; Garcia-Marin et al, 2019; Kelly et al, 2019) have demonstrated that in earlier studies there was a substantial underestimation of neuronal densities in different layers and sublayers of macaque V1 (O’Kusky and Colonnier, 1982; Beaulieu et al, 1992), For example, in the macaque, using the same techniques adopted in the current study, we reported that the overall neuronal density (Kelly and Hawken, 2017; Garcia-Marin., 2019; Kelly et al, 2019) was about double the density reported in earlier studies (O’Kusky and Colonnier, 1982; Beaulieu et al, 1992) but comparable to the density from a more recent study with extensive sampling using NeuN and stereological methods (Giannaris and Rosene, 2012).

The aim of the current study was to reevaluate the neuronal quantification in human V1 using large-scale imaging combined with automated image analysis and to make a systematic comparison between the human and macaque focusing on the laminar organization in terms of neuronal composition. While there have been numerous studies that have estimated the variation in neuronal density across layers in macaque, there are only a few studies in human (Inda et al., 2006; Leuba and Garey, 1989), and neither study measured continuously through the thickness of the cortex, from layer 1 (L1) to the white matter (WM). We found total cortical densities in human V1 that were 10-15% higher than those reported using stereological methods in Nissl-stained tissue (Everall et al, 1993, Pakkenberg and Gundersen, 1997; Dorf-Petersen et al, 2007). The human neuronal densities are about one third those we found in macaque V1. In addition, we determined the variation in neuronal density continuously through the thickness of cortex and found that there were substantial changes in neuronal density both between layers and within layers, like what we reported in macaque (Kelly and Hawken, 2017; Garcia-Marin et al, 2019).

Reconciling the differences in neuronal density with known allometric scaling in the Catarrhines, primates in general and across all mammals has major implications for understanding the relationship of structure to function (Finlay et al, 2001; Hill et al, 2010; Herculano-Houzel, 2014; Herculano-Houzel et al, 2015a,b). One means of relating the differences in density to other features of cortex is to define a module of cortex. Having an accurate estimation of the neuronal density allows the determination of the size of the populations underlying a processing module in cortex. One module that has been widely used in the comparative process was introduced by (Rockel et al, 1980). The proposed unit was the number of neurons under 1 mm^2^ of cortex through the full depth of cortex – from layer 1 to white matter. This module has been used in comparison of different mammalian cortices (Carlo and Stevens, 2013; Srinivasan et al, 2015) and within the Catarrhines (Colonnier & O’Kusky, 1981; O’Kusky & Colonnier, 1982) and across a wider range of primate species (Atapour et al, 2019).

However, as clearly pointed out by Rakic (2008) there are numerous types of columns or modules. A module that has been of interest since its discovery and elaboration is the ocular dominance column (ODC) (Hubel and Wiesel, 1962; 1968; Hubel et al, 1978; Horton and Hedley-Whyte, 1984; Adams et al, 2007) that is thought to contain the circuits for the emergence of orientation preference and spatial processing of a point image (Carlo and Stevens, 2013) in many mammalian species. In the current study we determined the population of neurons within each layer underlying an eye dominance module to compare with the size of the populations underlying the same module in macaque (Garcia-Marin et al, 2019) thereby evaluating the uniformity (Rockel et al, 1980; Carlo and Stevens, 2013) or nonuniformity (Herculano-Houzel et al, 2008; Lent et al, 2012) of cortex hypothesis at the level of the eye dominance module.

Although neuronal density and the size of ODCs and minicolumns are different in human and macaque, our current results show that there are a similar number of neurons in one minicolumn in both human and macaques. However, there are 1.7 times more minicolumns per ODC in human V1 compared to macaque V1 and therefore about twice the number of neurons. Although there are more ODCs in macaque than human in the macular region, there are more neurons in the macular ODCs in human than macaque and these human neurons are larger and have larger basal dendritic arbors than in macaque. Thus, our results suggest that for the same eccentricity, more neurons are devoted to the first visual processing steps in humans than in macaques, which, in turn, indicates the potential for a refined level of perceptual processing.

## MATERIAL AND METHODS

### Human brain tissue

Brain sections from seven adult humans were obtained from NIH NeuroBioBank. Specific information on each individual is presented in Supplementary Table 2. Briefly, all sections were from males without any reported neurological disease (mean age, 51.7 ± 5.2 y; range, 23 to 66 y). After death, the left hemisphere was fixed in 4% PFA, and 40 µm thick coronal cryostat sections were obtained from regions of primary visual cortex (V1) that had a visual field representation of between 2.5 – 5 deg. eccentric at the floor of the calcarine sulcus – near to the horizontal meridian.

### Immunofluorescence

To estimate the total neuronal density and the density of the PV inhibitory neurons, sections were immunofluorescence stained for the pan-neuronal marker NeuN and for the calcium-binding protein parvalbumin (PV) (Fig. 1A-B). Sections were incubated for 1h in a blocking solution of 0.01 M PBS with 0.25% Triton-X and 3% normal goat serum (NGS) and then incubated overnight at 4 °C with rabbit anti-NeuN antibody (1:2000, MAB5504, Millipore, Temecula, CA, USA) in 0.01 M PBS with 0.25% Triton-X and 3% NGS. Sections were rinsed in 0.01 PBS and incubated overnight at 4 °C with Alexa biotinylated goat anti-rabbit 488 (1:2000, Vector Laboratories, Burlingame, CA). After rinses, sections were incubated overnight at 4°C in guinea pig anti-PV polyclonal serum (1:1000; 195 004, Synaptic Systems, Göttingen, Germany), and later in Alexa biotinylated goat anti-guinea pig 594 (1:500; BA-7000, Vector, Burlingame, CA, USA). After rinsing in PBS the sections were counterstained with 10 µg/mL DAPI (D9542-5MG, Sigma–Aldrich, St. Louis, MO, USA; Fig. 1C) and were treated with Autofluorescence Eliminator Reagent (2160, Millipore) to minimize lipofuscin-like autofluorescence. Finally, the sections were washed and mounted with ProLong Gold Antifade Reagent (Invitrogen Corporation, Carlsbad, CA, USA).

**Figure 1:**
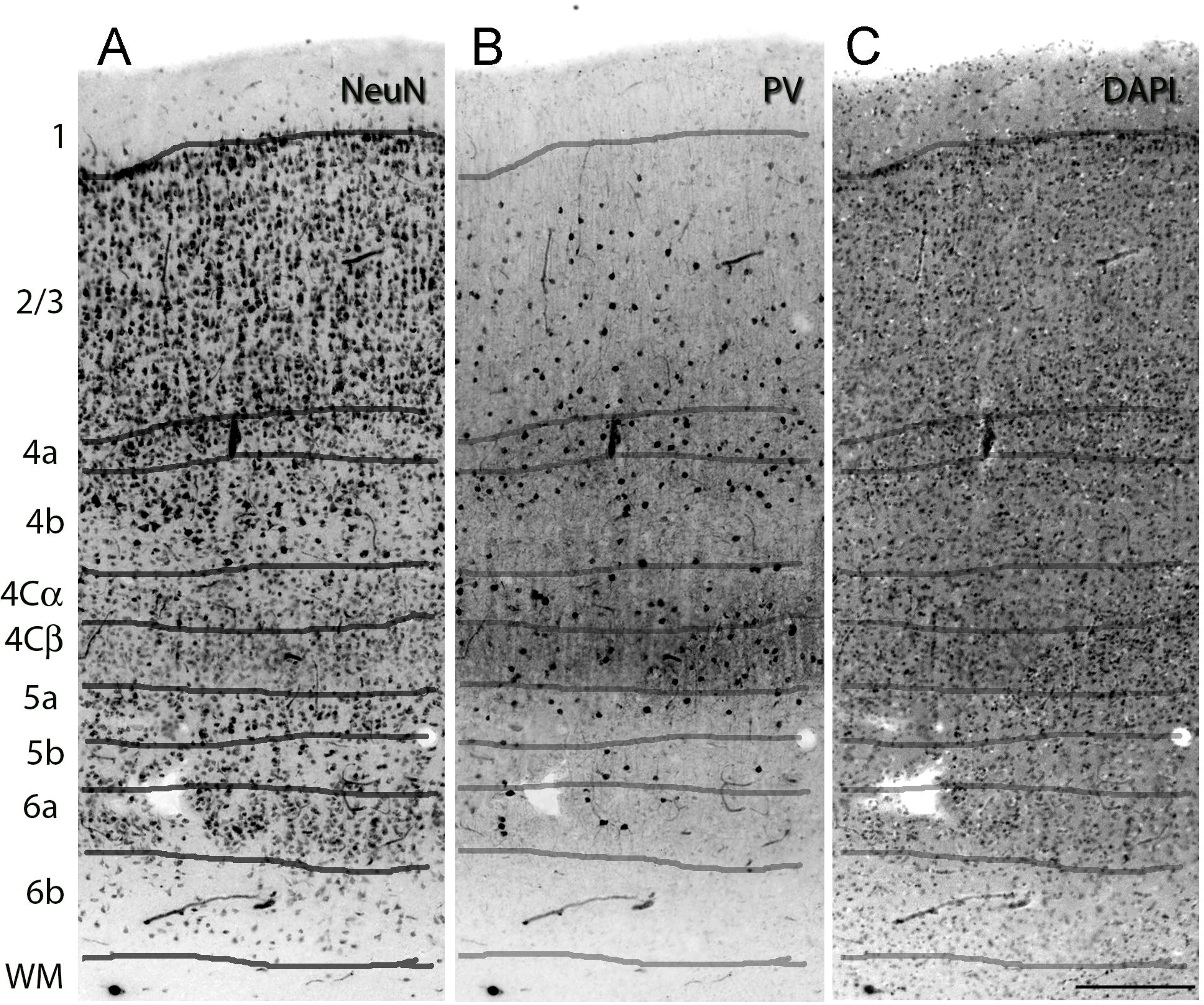
Laminar distribution of neurons through the thickness of human V1. Section was triple labeled for NeuN, PV and DAPI. The laminar boundaries are drawn according to the scheme of (Brodmann 1909; Braak, 1982). (A), neuronal population immunoreacted with the pan neuronal marker NeuN. (B), neurons in V1 immunoreactive for parvalbumin (PV). (C) the total cellular population including neurons, glial and epithelial cells labelled with DAPI. Scale bar in C 250 μm.

### Calculation of the neuronal density

For neuronal quantification, fluorescence sections were imaged using a Leica TCS SP8 confocal system (Leica Microsystems, Wetzlar, Germany). A 488 nm Argon laser, a 564 nm DPSS laser, and a 405 nm diode laser were used to excite the Alexa Fluor 488, and 594 and DAPI, respectively. Gain and offset levels were set for each channel such that there were few saturated pixels and minimal background noise. Image stacks were acquired by specifying an upper and lower z position, which correspond to the top and the bottom of the section, as we were able to get complete penetration of the antibodies through the 40 µm section.

For quantification of neuronal density, we followed the methodology previously described in our recent studies (Kelly and Hawken, 2017; Kelly et al, 2019; Garcia-Marin et al, 2019). Briefly, a column of tissue was defined spanning from the pial surface to the white matter and oriented orthogonal to the pial surface (Fig. 2A). Two columns were acquired for each section from each subject. The Leica Application Suite X software automatically adjusted the number of stacks needed to cover the selected region; in our case this number ranged from 9 to 14 stacks, with constant horizontal overlap of 110 pixels between stacks. Each stack was acquired using a 40X oil-immersion objective lens (NA 1.4, refraction index 1.45), zoom 1, with a pinhole size of 1 airy unit, z-step of 0.5 µm, a range of 54-103 (mean 74) optical planes. The z stack acquisition routinely included planes above and below the section surfaces. The image resolution in x-y was 1024x1024 pixels (290.6 x 290.6 µm), and the scanning speed was 400 Hz.

**Figure 2:**
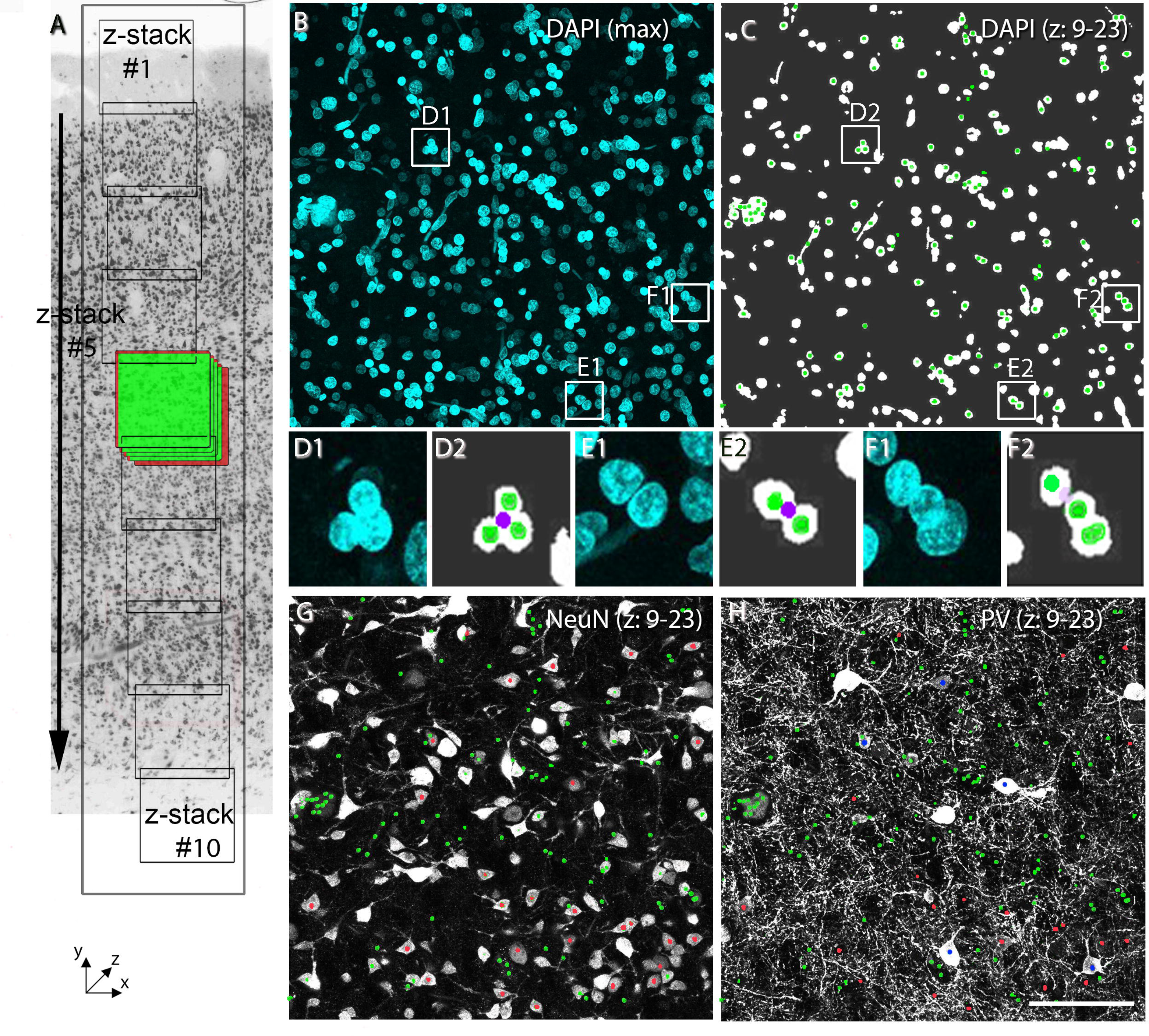
Methods for quantifying cellular (DAPI), neuronal (NeuN) and parvalbumin (PV) densities through the different cortical layers. (A) A column of tissue spanning from the pial surface to the white matter and oriented orthogonal to the pial surface. A vertical series of z-stacks, spanning the thickness of cortex (each stack: 1024x2014 pixels, 290.6 x 290.6 µm, 40X oil-immersion objective lens, zoom 1, z-step of 0.5 µm) was acquired. The number of z-stacks required to span the thickness varied between 9 and 14, depending on the individual section, for example in (A) 10 z-stacks were acquired; z-stack#1 was acquired at the pial surface and z-stack#10 at the layer 6/WM boundary. Within each z-stack, a range of 54-103 optical planes were acquired to span the 40 μm thick section, example indicated for the blue z-stack #5 in (A). For the neuronal quantification, within each z-stack we defined inclusion and exclusion borders of the counting frame, green (right, down, and top) and red borders (left, up, bottom), respectively around z-stack 5. (B) Maximum projection of 70 images from the DAPI channel. Three groups of cells are shown at higher magnification in the white squares (D1, E1, F1). (C) Image of a maximum projection of 14 images from the DAPI channel. (D1-D2), (E1-E2), and (F1-F2) inserts from B and C, respectively, showing the cluster of neurons in each white square (D1, E1, and F1) and how they are individually resolved in the segmentation process (D2, E2, and F2). Each purple dot shows the centroid of the cluster and each green dot shows the centroid of the individual cells in each cluster extracted by the automated segmentation algorithm. (G) Image of a maximum projection of 14 images from the NeuN channel, in which DAPI centroids are labeled with a green dot and the DAPI centroids that colocalize with a neuron are identified with a red dot. Note that the green (DAPI) centroids that do not colocalize with a NeuN-ir neuron belong to a non-neuronal cell. The NeuN-ir neurons that do not colocalize in this subset of z-planes have their centroid in other image planes. (H) Image of a maximum projection of 14 images from the PV channel, in which DAPI centroids are labeled with a green dot and the DAPI centroids that colocalize with NeuN and PV are identified with a blue dot. The red dots indicate the NeuN labeled neurons shown in G. Note, some PV-ir neurons that have neither DAPI or NeuN centroids have their centroid in other image planes. Scale bar: 290 µm (A),85 µm (B, C, G, H), 20 µm (D1-F1 and D2-F2).

Next, DAPI, NeuN and PV images were automatically analyzed, and each cell was categorized according to its pattern of immunoreactivity, using the method previously described (Kelly and Hawken, 2017; Kelly et al, 2019) (Fig. 2B-H). Briefly, cell centroids are identified in 2D for each optical plane in the DAPI channel (Fig. 2B-C). In a single image plane the DAPI-labeled cells could be in close proximity to each other, so the 2D segmentation provided one single centroid for a cluster of cells (Fig. 2D1-F2; magenta dots). The next step identified the 3D centroids of the cells and split the clump into individual cells (Fig. 2D1-F2; green dots). Once the 3D centroids were identified for each cell, the other channels were evaluated for marker expression at the locations of the centroids (Fig. 2G-H). Cells were quantified using stereological exclusion boundaries (Sterio 1984; Gundersen et al. 1988) where each stack was treated as a 3D counting brick probe (Howard and Reed 20055; Williams and Rakic 1988). Objects touching the top, left or back planes were excluded from quantification, whereas objects touching the bottom, right or front planes were included.

Finally, the neuronal counts were converted to densities by dividing the number of counted neurons by the sampled volume in each stack, corrected to account for tissue shrinkage. The sample volume in each stack was determined as the region in which the penetration of the antibodies was uniform. Although the antibodies penetrated through the whole depth of the sections, optical planes acquired near the cut tissue surfaces yielded lower counts than optical planes acquired in the middle of the section. For each z stack in the column, we determined the z-range where the counts plateaued, then found the minimum plateau-range across all the z stacks and applied this range to all the stacks in the same column. On average we sampled 20±3 optical planes (Table 1). To determine the laminar density, laminar boundaries in V1 were identified in each column by visually identifying transitions in cell density and neuronal composition (Fig. 1) consistent with previous descriptions of these features in V1 (Brodmann 1909; Braak, 1982)

Using the same z-stacks we also calculated the volume occupied by the soma and proximal dendrites of the neurons in the images labeled with NeuN. For each stack, we applied a Gaussian filter of 2.0, automatically threshold the images, and obtained the volume occupied for the neurons in the neuropil using the tool ‘Analyze Particles’ of ImageJ.

### Calculation of Shrinkage

The degree of tissue shrinkage was measured by comparing sections in x, y, and z dimensions before and after tissue preparation with the aid of Olympus VS120 and confocal Leica SPC8. The average linear shrinkage (S) was calculated by using the following formula: S = (A−0)/A, where A is the absolute value before processing and 0 is the observed valued after processing. This yielded an average linear shrinkage of 18%, 18%, and 20%, in x, y, and z, respectively.

### Image Processing for Figures

Images presented in figures were captured using a Leica TCS SP8 confocal system, or an Olympus VS120-FL virtual slide scanning system. ImageJ was used to generate maximum intensity z projections. Adobe Photoshop CS software (Adobe Systems, San Jose, CA) was used to adjust the images for brightness and contrast, and to generate the figure plates. Images were not otherwise altered in any way, e.g., by removing or adding image details.

## RESULTS

Initially we show the neuronal laminar density distribution in human V1 and the laminar density distribution of the largest population of interneurons, the PV neurons. Next, we estimate the number of neurons in each layer through the depth of an ocular dominance processing module in human V1 and make a comparison to the same processing unit in macaque monkey visual cortex. The results indicate that although the density of neurons (per mm^3^) is considerably lower in humans than macaque the number, when normalized to the same cortical processing module, is higher in humans than macaque.

### Neuronal and glial density using 3D-confocal imaging

Many previous studies have estimated the neuronal density in human V1 using non-stereological techniques in 2D and reported a wide range of average density values. We recently demonstrated (Kelly and Hawken, 2017; Garcia-Marin et al, 2019) that these techniques produced a substantial underestimation of the total neuronal density. Average neuronal density values estimated using stereology are higher than those reported from non-stereological studies but also show substantial variation between studies (40.5 – 102 x 10^3^ neurons/mm^3^).

We estimated the total neuronal density using NeuN and the total density of the largest subpopulation of interneurons, the PV neurons. Overall, we found a neuronal density of 79.2 ± 3.0 x 10^3^ neurons/mm^3^ (mean ± 1 sem; Table 1, n = 13 from 7 individuals). We found a DAPI density of 164.9 ± 5.8 x 10^3^ cells/mm^3^ and a total PV density of 4.6 ± 0.4 x 10^3^ PV-ir neurons/mm^3^, which represents 5.8% of the total number of neurons. The mean vertical cortical thickness (pia to white matter) was 2.8 ± 0.1 mm.

A total of 12,256 neurons were counted in a total volume of 0.146 mm^3^, with an average volume of 0.012 mm^3^ per individual. For six individuals (H2-H6) the mean neuronal density across all layers showed a relatively narrow range (74,840-90,880 neurons/mm^3^). However, for one individual (H1) the mean neuronal density was low compared with the other cases (63,720 neurons/mm^3^) although this difference is not significant. The mean pairwise difference between measurements within an individual was about 10% of the mean while the average between subject difference was 16% of the mean. These two distributions were not significantly different (Student’s ttest, p = 0.25). The direction of this result is in agreement with previous observations that variation in neuronal density between individuals of similar ages exceeds the variation in repeated measures within an individual (Leuba and Garey, 1989). Furthermore, there was no correlation between the neuronal density and age (r^2^ = 0.04126, ns) (Suppl. Fig. 2).

#### Glia to neuron ratio

DAPI labels not only neurons but also all other cell types, which in the cortex primarily consist of glial cells and endothelial cells. The density of non-neuronal cells is the total density of DAPI-labeled cells (165 x 10^3^ cells/mm^3^; Table 1) minus the neuronal density (79 x 10^3^ neurons/mm^3^; Table 1), resulting in a density of non-neuronal cells of 86 x 10^3^ cells/mm^3^. Using a prior estimate that endothelial cells make up about 30% of the total non-neuronal cells (von Bartheld et al, 2016), we estimated that about 70% of the non-neuronal population were glial cells (60 x 10^3^ cells/mm^3^). Therefore, we determined that the average glia to neuron ratio (60/79) was 0.76 and the non-neuronal cell to neuron ratio (86/79) was 1.1.

### Laminar Distribution of Neuronal Density

The density was determined continuously, from sequentially acquired images, from the pial surface to the white matter (a column through the thickness of cortex) and this allowed us to analyze the neuronal distribution within layers (Fig. 3A) and across the normalized cortical thickness (Fig. 3B). Neuronal density varies at a finer scale than the divisions between the major cortical layers. Because of the different relative position of the laminar boundaries in each individual section and even between columns within a section and the fine scale density variation, we adopted an alignment strategy where we found the peaks and the valleys in each individual column. The peaks correspond to the position of the highest density in layer 2, layer 3B/4A, layer 4Cβ and layer 6; with the valleys the positions in layer 4B and the border 4Cβ/5. Using this strategy, we aligned the different columns and show the result in Figure 3B and Table 3. Below we present the results based on conventional layer boundaries and then compare these results to those determined by aligning peaks and valleys between columns.

**Figure 3:**
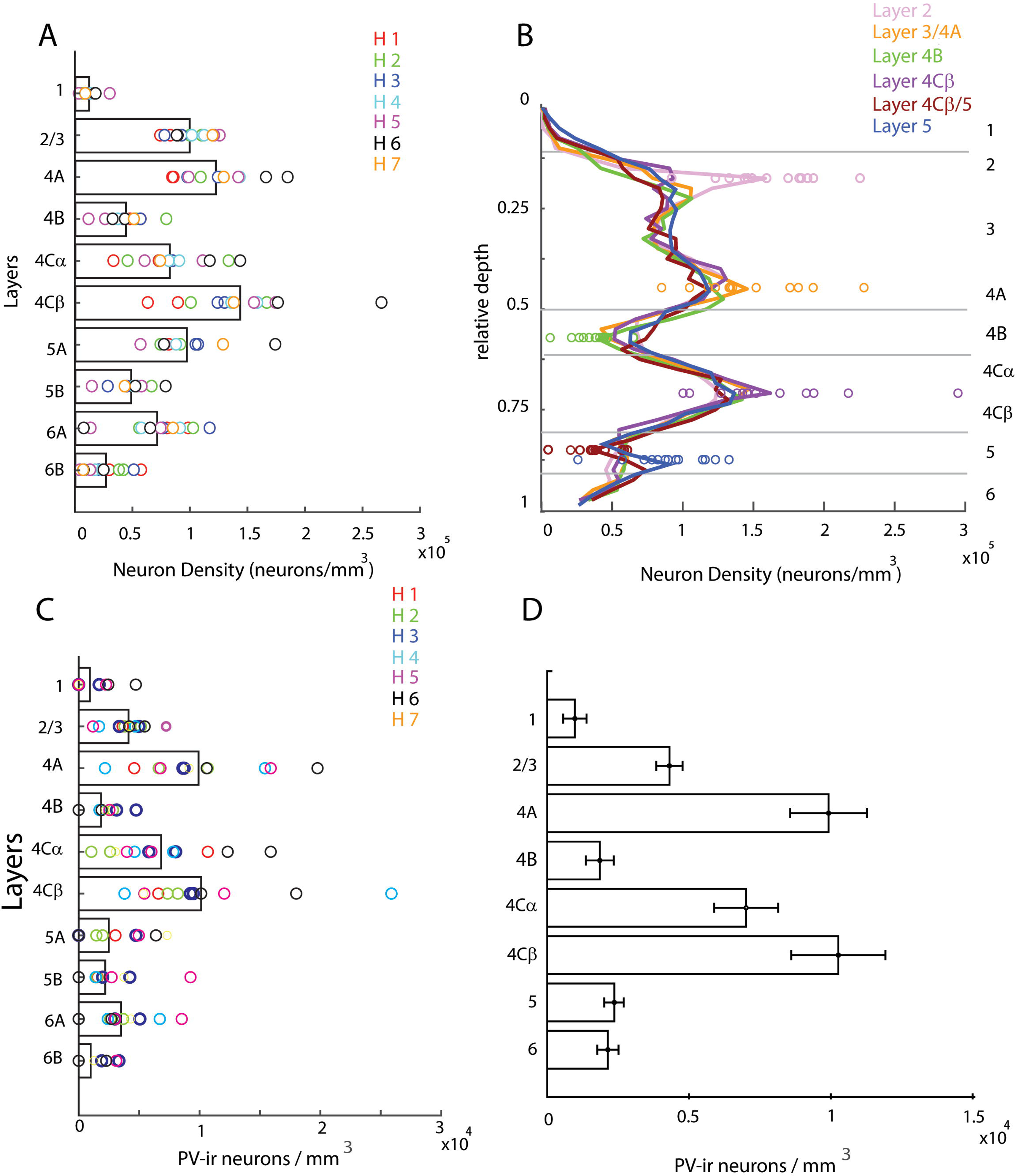
(A) Laminar density of neurons in human V1. Average density for 13 columns from 7 individuals for each layer shown by the horizontal bars. Individual layer density values for each column are represented by ‘o’. The values from each individual (H1 – H7) are shown by the colors in the legend. There are two columns from each individual except H7 where there is one. (B) Average continuous density through the relative cortical depth. The cortical depth was normalized for each section. The densities were individually smoothed for each section. Next the density distributions were aligned to a peak – layer 2 (pink), border 3/4A (orange), layer 4Cβ (purple) and layer 5 (blue) – or valley – layer 4B (green) and 4Cβ/5 (brown) – and averaged. The values from each section sampled at the aligned peak or valley are shown by the open circles. C) Laminar density of parvalbumin-ir (PV) neurons per mm^3^ in human V1. Individual values for each column are represented by ‘o’. D) Average density of PV neurons per mm^3^ is shown by the height of the individual bars, the error bar indicates the SEM. Average density determined for 13 columns from 7 individuals.

**Table 2:** Human neuronal density by layers (neurons/mm^3^) for each of the 7 human (H1-H7) and each sampling columns (b1-b2)

| layers<br>human | 1 | 2/3 | 4a | 4b | 4Cα | 4Cβ | 5a | 5b | 6a | 6b |
| --- | --- | --- | --- | --- | --- | --- | --- | --- | --- | --- |
| H1b1 | 9,452 | 82,280 | 85,037 | 56,363 | 32,710 | 88,841 | 80,877 | 52,956 | 97,829 | 56,751 |
| H1b2 | 5,770 | 73,387 | 83,880 | 46,950 | 72,110 | 62,584 | 80,574 | 53,330 | 79,442 | 29,064 |
| H2b1 | 5,929 | 99,198 | 125,100 | 52,176 | 45,051 | 100,085 | 90,912 | 43,480 | 55,115 | 37,483 |
| H2b2 | 9,970 | 116,903 | 117,293 | 78,631 | 88,500 | 160,004 | 124,186 | 60,389 | 111,051 | 59,944 |
| H3b1 | 11,862 | 92,077 | 97,066 | 56,237 | 84,266 | 129,453 | 106,336 | 44,439 | 116,464 | 50,432 |
| H3b2 | 10,909 | 77,154 | 123,653 | 41,409 | 73,163 | 123,365 | 104,200 | 27,719 | 75,860 | 19,991 |
| H4b1 | 13,420 | 111,650 | 97,927 | 43,135 | 80,903 | 135,321 | 87,177 | 54,771 | 90,583 | 16,775 |
| H4b2 | 13,396 | 100,959 | 142,620 | 36,454 | 89,931 | 157,873 | 76,429 | 50,726 | 57,034 | 22,792 |
| H5b1 | 29,439 | 125,158 | 98,125 | 11,008 | 59,834 | 173,919 | 56,268 | 13,877 | 12,788 | 3,450 |
| H5b2 | 2,571 | 120,349 | 141,436 | 25,427 | 110,406 | 155,070 | 77,109 | 57,210 | 73,771 | 12,437 |
| H6b2 | 16,541 | 88,651 | 165,519 | 42,720 | 116,544 | 175,857 | 76,729 | 51,762 | 6,794 | 24,155 |
| H6b3 | 17,137 | 87,863 | 184,344 | 31,970 | 143,025 | 266,047 | 173,617 | 78,150 | 64,702 | 6,836 |
| H7b2 | 8,150 | 118,999 | 128,712 | 50,507 | 73,349 | 137,485 | 128,097 | 42,529 | 84,496 | 6,334 |
| **Mean** | **11,888**<br>**(16%)** | **99,587**<br>**(133%)** | **122,362**<br>**(164%)** | **44,076**<br>**(59%)** | **82,292**<br>**(115%)** | **143,531**<br>**(194%)** | **97,116**<br>**(125%)** | **48,564**<br>**(66%)** | **71,225**<br>**(95%)** | **26,650**<br>**(34%)** |
| **SEM** | 2,746 | 5,036 | 9,858 | 3,954 | 8,720 | 15,163 | 9,355 | 5,954 | 11,206 | 6,884 |

**Table 3:** Human neuronal density (neurons/mm^3^) when peaks or valleys are aligned for each of the 7 human (H1-H7) and each sampling columns (b1-b2).

| layers<br>human | 2 | 3/4a | 4b | 4Cβ | 4Cβ /5 | 5 |
| --- | --- | --- | --- | --- | --- | --- |
| H1b1 | 144,620 | 105,680 | 38,935 | 105,680 | 38,935 | 116,810 |
| H1b2 | 133,805 | 118,940 | 47,575 | 101,100 | 35,681 | 83,260 |
| H2b1 | 160,258 | 133,990 | 42,035 | 139,240 | 36,780 | 73,560 |
| H2b2 | 146,830 | 136,190 | 65,970 | 193,640 | 61,710 | 114,900 |
| H3b1 | 124,016 | 124,020 | 44,646 | 146,340 | 59,528 | 124,020 |
| H3b2 | 93,017 | 134,870 | 44,183 | 127,900 | 27,905 | 79,060 |
| H4b1 | 183,290 | 133,650 | 30,548 | 152,740 | 45,822 | 133,650 |
| H4b2 | 189,269 | 182,740 | 39,159 | 169,690 | 39,159 | 97,900 |
| H5b1 | 193,376 | 85,940 | 7,162 | 143,240 | 21,486 | 21,486 |
| H5b2 | 149,912 | 176,820 | 34,595 | 188,350 | 57,659 | 80,720 |
| H6b2 | 226,371 | 193,240 | 27,606 | 218,090 | 5,521 | 57,970 |
| H6b3 | 184,820 | 229,170 | 22,178 | 295,700 | 36,963 | 96,100 |
| H7b2 | 175,487 | 152,840 | 45,287 | 147,180 | 28,304 | 90,570 |
| Mean | **161,929** | **146,776** | **37,683** | **163,761** | **38,112** | **90,001** |
| SEM | 10,438 | 11,222 | 4,265 | 14,706 | 4,772 | 9,541 |

#### Supragranular layers

In the supragranular layers of cortex, layer 1 had a significantly lower density than all the other layers, 12 x 10^3^ neurons/mm^3^ (One-way Anova, Suppl. Table 3). The low density in layer 1 is a characteristic of this layer in all mammals (Marin-Padilla and Marin-Padilla, 1982, Gabbott and Somogyi, 1986, Balaram and Kaas, 2014). It is common to assign layers 2 and 3 together in density estimates. When this was done the mean density across individuals was 100 x10^3^ neurons/mm^3^, which was significantly lower than layer 4Cβ and higher than layers 4B, 5B, and 6B (Fig. 3A; Table 2; One-way Anova, Suppl. Table 3). However, within layers 2/3 there are quite consistent variations in density. In all the cases studied there was a local density peak in layer 2 (Fig. 3B, pink trace) followed by a trough and then another peak in lower layer 3 near the layer 3/4A border (Fig. 3B, orange trace). The local density in layer 2, when all the continuous estimates were aligned by the layer 2 peaks (Fig. 3B), reached a mean value of 162 x 10^3^ neurons/mm^3^ (Table 3). This suggests that there is sublayer organization that is not captured by the laminar density estimates. A second peak in lower layer 3 – probably including layer 4A – had a density of 147 x 10^3^ neurons/mm^3^ when all the columns were individually aligned (Fig. 3B, orange trace; Suppl. Table 3).

#### Granular layers

The layer with the highest mean laminar density was layer 4Cβ with 144 x 10^3^ neurons/mm^3^ (Fig. 3A), which was almost twice the average density across the whole cortex. This density was significantly higher than all layers except for layer 4A (Table 2, Suppl Table 3). There was also considerable variability between individuals (Fig. 3A); H6 had an average density of 221 x 10^3^ neurons/mm^3^ in layer 4Cβ whereas H1 had an average density of 76 x 10^3^ neurons/mm^3^ (Fig. 3A; Table 2). The within individual values were considerably less variable.

The other main TC recipient layer, 4Cα, where the main excitatory cell type is also the spiny stellate cell, had an average density of 82 x 10^3^ neurons/mm^3^ that was significantly different from layers 1, 4A, 4B and 6B (Figure 3A, Table 2, Suppl. Table 3). The density ratio between 4Cα and 4Cβ was 0.60. This ratio was remarkably consistent across individuals ranging from 0.52 to 0.69. When the densities were aligned to the peak in 4Cβ the mean density at the peak depth bin was 164 x 10^3^ neurons/mm^3^ (Fig, 3B, purple trace; Table 3).

The precise boundaries of layer 4A are difficult to define in human visual cortex because there are no distinct upper or lower layer boundaries such as those provided by the honeycomb arrangement of CO in Old-World monkey V1 (Horton, 1984; Garcia-Marin et al, 2013) and vGlut2 (Garcia-Marin et al, 2015) that provides clear boundaries. We defined layer 4A at low power as a thin strip of high neuronal density, below layer 3. Using these criteria to determine the layer 4A boundaries our best estimate of the layer 4A density was 122 x 10^3^ neurons/mm^3^ (Fig. 3A, B, Table 2). The neuronal density in this layer was significantly higher than in layers 4B, 4Cα, 5B, 6A-6B. Finally, layer 4B had an average density 44 x 10^3^ neurons/mm^3^ (Fig. 3A), and when the 4B troughs were aligned the density was reduced to 38 x 10^3^ neurons/mm^3^ (Fig, 3B, green trace; Table 3.).

#### Infragranular layers

The infragranular layers 5 and 6 occupy 30% of the vertical extent of cortex (Table 5), and account for about 22% of the total neuronal population (Table 5). The highest densities are in the upper halves of layers 5 and 6 (97 x 10^3^ neurons/mm^3^ and 71 x 10^3^ neurons/mm^3^ respectively (Table 2). The neuronal density in layer 5A was significantly higher than in layers 5B and 6B. The average density in layer 6B (27 x 10^3^ neurons/mm^3^, Table 2) was lower than in layer 5B, and significantly lower in layer 6B than in 6A (Suppl. Table 3). There is a consistent valley in layer 5 and when these valleys were aligned, the density at the trough was 38 x 10^3^ neurons/mm^3^ (Fig. 3B, brown trace; Table 3). Finally, there is a local peak at the layer 5/6 border (Fig. 3B, blue trace; Table 3) where the density rises to 90 x 10^3^ neurons/mm^3^.

**Table 4:**
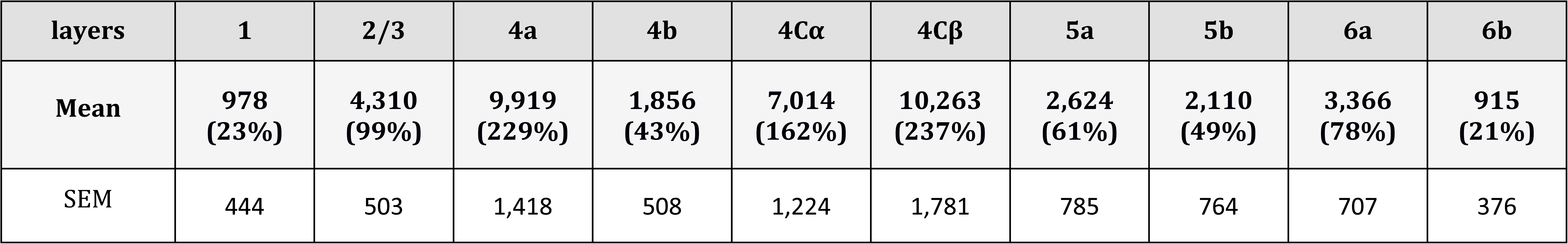
Mean PV density (neurons/mm^3^) by layers across all human (H1-H7) and columns (b1-b2), and the percentage of PV in each layer with respect to the mean PV density across all layers. (Mean PV density across all layers 4,605 neurons/mm^3^ (see table 1)).

**Table 5:** Calculations of the total number of neurons, excitatory, and inhibitory neurons per ocular dominance column (ODC) in human by layers.

| Layer | layer width (mm) | density (neu/mm<sup>3</sup>) | neurons/ODC (0.86*0.86 * width) | GABA % | excitatory neurons/ODC | inhibitory neurons/ODC |
| --- | --- | --- | --- | --- | --- | --- |
| 1 | 0.281 | 11,888 | 2,471 | 67 | 815 | 1,655 |
| 2/3 | 0.722 | 99,587 | 53,178 | 13 | 46,265 | 6,913 |
| 4a | 0.208 | 122,362 | 18,914 | 12 | 16,645 | 2,270 |
| 4b | 0.301 | 44,076 | 9,812 | 12 | 8,635 | 1,177 |
| 4Cα | 0.205 | 82,292 | 12,477 | 13 | 10,855 | 1,622 |
| 4Cβ | 0.192 | 143,531 | 20,382 | 11 | 18,140 | 2,242 |
| 5a | 0.163 | 97,116 | 11,708 | 8 | 16,670 | 1,450 |
| 5b | 0.162 | 48,564 | 6,142 |  |  |  |
| 6a | 0.238 | 71,225 | 12,063 | 6 | 16,000 | 1021 |
| 6b | 0.238 | 26,650 | 4,691 |  |  |  |
|  | **2.7** | **79,221** | **158,843** | **11** | **141,370** | **17,473** |

#### Comparison with layer distribution in macaque

Recently we studied the neuronal laminar density in macaque V1 (Kelly et al, 2019), using the same methodology as in the current study. To compare the laminar densities between macaque and human V1 we used the mean density across all layers to normalize the layer data and then compared the densities within layers as deviations around the mean (Fig. 4A). Although the absolute density in macaque V1 is about 3 times greater than the density in human V1 the relative distribution across layers within cortex is remarkably similar between macaque and human (Fig. 4A).

**Figure 4:**
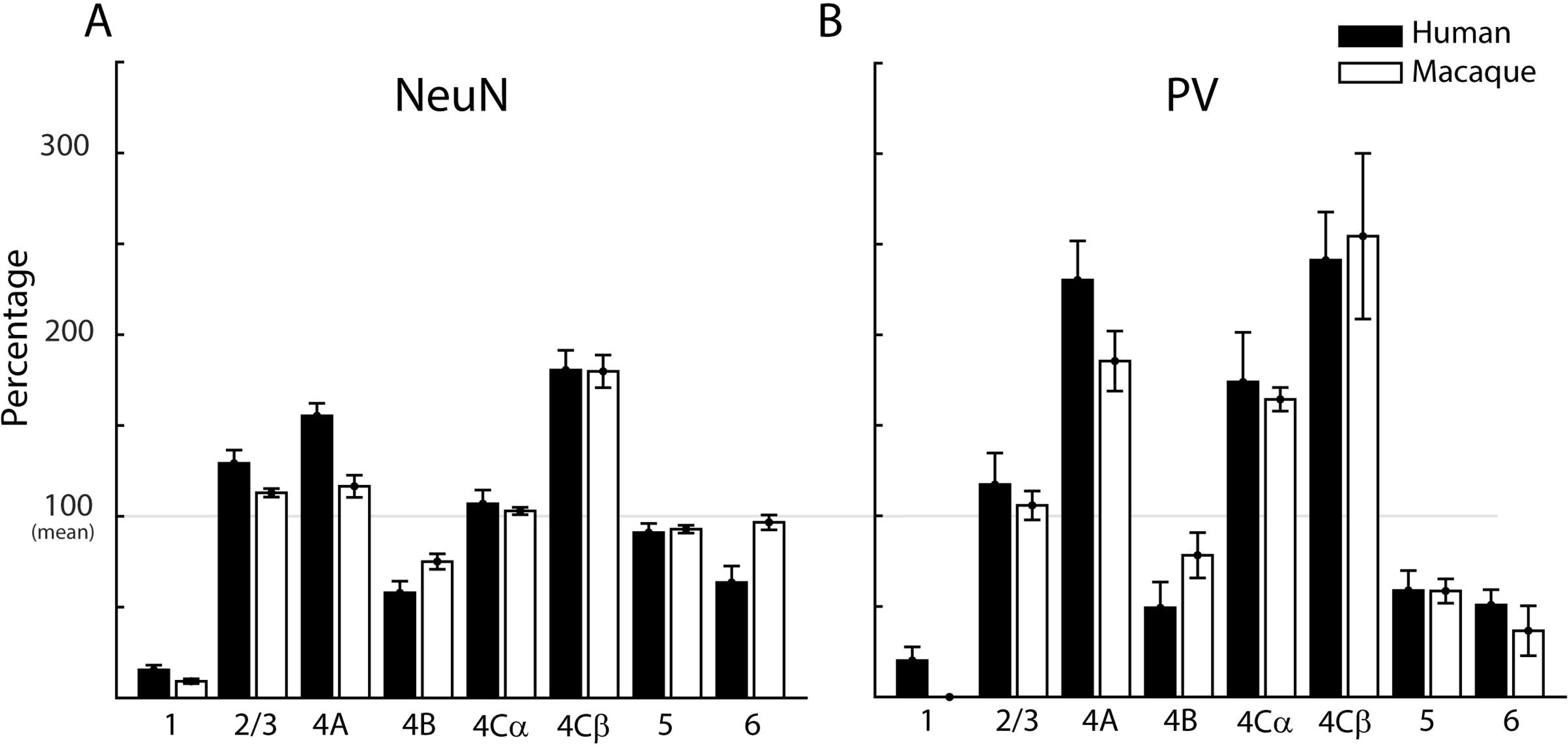
Comparison of the laminar distribution of neurons between human (green) and macaque (yellow). (A) Relative percentage of Neun-ir neurons in each layer. (B) Relative percentage of parvalbumin neurons (PV-ir). The values shown for each layer were determined as a percentage relative to the total average cortical density (shown for visualization as the grey horizontal line at 100%). Macaque data from Garcia-Marin et al., 2019, Kelly et al., 2019. Error bars indicate the SEM.

Using the same z-stacks we calculated the volume of neuropil occupied by neuronal somata and proximal dendrites in the NeuN sections using ImageJ. In human, neurons occupied 7.5% of the neuropil, on average, across all the layers. Using the data from macaque, Kelly and Hawken (2017), we also calculated the volume of neuropil occupied by the neuronal cell bodies and proximal dendrites in the NeuN sections across all layers. In macaque the soma of the neurons and their proximal dendrites occupied 15.5%, suggesting that there is more neuropil volume in human than macaque that could be occupied by other elements rather than soma and proximal dendrites of neurons.

### Distribution of PV Neurons

PV-ir interneurons comprise the largest population of inhibitory interneurons in cortex (Rudy et al, 2011). Previously, in macaque V1, we found that the population of GABAergic interneurons was about 11% of the total population and PV-ir neurons were ∼ 52% of the GABAergic population, therefore 5-6% of the total population (Kelly et al, 2019). In the current study we found a total density of 4.6 x 10^3^ PV neurons/mm^3^, which represents 5.8% of the total neuronal population (Table 1), similar to the percentage that we previously reported for macaque (Kelly et al, 2019). We observed the highest density of PV neurons in 4Cβ with 10.3 x 10^3^ neurons/mm^3^ followed by layer 4A with 9.9 x 10^3^ neurons/mm^3^ and layer 4Cα with 7.0 x 10^3^ neurons/mm^3^ (Fig. 3C-D; Table 4). Layers 2/3, 6A, 5A, 5B have lower densities ranging from 2 - 4 x 10^3^ neurons/mm^3^. The lowest densities were found in layers 1 and 6B, with, 1.0 and 0.9 x 10^3^ PV neurons/mm^3^, respectively.

When we compared the normalized densities across layers in human to those in macaque V1 (Kelly et al, 2019) we found the PV neurons within each layer – as a proportion of the mean density across all layers – was remarkably similar between the human and the macaque (Fig. 4B). A similar close correspondence was found for the total neuron density (Fig. 4A). However, there is nearly a two-fold difference in the proportion of neurons that are PV-ir in the putative output layers 2/3, 4B, 5A, 5B, 6A and 6B, where about 4% of neurons are PV-ir, compared to input layers 4A, 4Cα and 4Cβ, where more than 8% of neurons are PV-ir.

### Connectivity

We next sought to compare not just differences in density between humans and macaque monkeys but the composition of comparable processing units. The cortex is often thought of as a collection of repeating columnar structures. At the finest scale the processing unit is the cortical microcolumn (Mountcastle et al, 1957; Mountcastle, 1997). In primary visual cortex, at a meso-scale the cortical ocular dominance column (ODC), structurally demarcated by cytochrome oxidase patches (Horton and Hedley-White, 1984; Adams et al, 2007), is considered an important processing unit (Hubel and Wiesel, 1977; Lund et al, 2003) that contains representations of the full range of orientations at a range of spatial scales for one eye.

Adams et al (2007) found that ODC widths in human V1 were, on average, 863 μm. If it is assumed that each eye dominance patch defines a column (Hubel et al, 1. 1978) then, in human cortex where the vertical extent (pia to white matter) is on average 2700 μm, an ODC volume is 863 μm x 863 μm x 2700 μm. In the cynomolgus macaque monkey the ODC volume is 530 μm x 530 μm (Horton and Hocking, 1996) x 1500 μm. The ratio of the ODC area tangential to the cortical surface is 2.6:1 (human:macaque) and the ratio of ODC volumes is 4.8:1 (human:macaque).

There is some debate as whether each ODC has the same dimensions as an orientation domain (see Mazade and Alonso, 2017 for summary across species). Nonetheless, the human/macaque ratio of surface area is likely to be similar for orientation domains and ODCs, and the difference in thickness of cortex between the human and macaque is well established (Wagstyl et al, 2015; Alvarez et al, 2019). In what follows we will use the ODC to compare between human and macaque.

We estimated the total number of neurons under one ODC (Table 5). Using the total neuronal density calculated in the current study (Table 1), we determined there were 159 x 10^3^ neurons in one ODC (Table 5, see Table 7 for calculations) in human V1. The same calculation applied to our previous macaque data (Garcia-Marin et al, 2019) showed there were 96 x 10^3^ neurons in one ODC. Consequently, there are 1.7 more neurons in an ODC in human V1 than in macaque. Although there was a considerable range in the ratio of the number of neurons in an ODC when comparing within layers between human and macaque, two factors were of considerable interest. The principal thalamic recipient layers (4Cα and 4Cβ) had about the same number of neurons in an ODC in human as in macaque. In contrast, the output layers had a greater number of neurons in the human ODC when compared to macaque. Because we could not easily distinguish a layer 4A/4B boundary in human V1 we summed the neurons in these two layers (4A and 4B) and compared the number to the sum from the macaque; there were 2.4 times the number of neurons in the human ODC. Layers 2/3 and 5 had 1.7 and 1.9 times more neurons in a human ODC. Layer 6 had 1.2 times the number. Layer 1, which has considerably greater thickness in human than in macaque has 3.8 times more neurons in the human ODC.

**Table 6:** Calculations of the total number of neurons, excitatory, and inhibitory neurons per ocular dominance column (ODC) in monkey by layers (from Garcia-Marin et al., 2019)

| Layer | layer width (mm) | density (neu/mm<sup>3</sup>) | neurons/ODC (0.53*0.53 * width) | GABA % | excitatory neurons/ODC | inhibitory neurons/ODC |
| --- | --- | --- | --- | --- | --- | --- |
| 1 | 0.112 | 20,700 | 651 | 67 | 215 | 436 |
| 2/3 | 0.434 | 257,800 | 31,429 | 13 | 27,343 | 4,086 |
| 4a | 0.056 | 268,800 | 4,228 | 12 | 3,721 | 507 |
| 4b | 0.154 | 173,800 | 7,518 | 12 | 6,616 | 902 |
| 4Cα | 0.172 | 235,000 | 11,368 | 13 | 9,891 | 1,478 |
| 4Cβ | 0.172 | 411,000 | 19,838 | 11 | 17,656 | 2,182 |
| 5 | 0.154 | 213,000 | 9,214 | 8 | 8,477 | 737 |
| 6 | 0.238 | 214,500 | 14,340 | 6 | 13,480 | 860 |
|  | **1.5** | **229,100** | **96,016** | **11** | **85,455** | **10,562** |

**Table 7:**
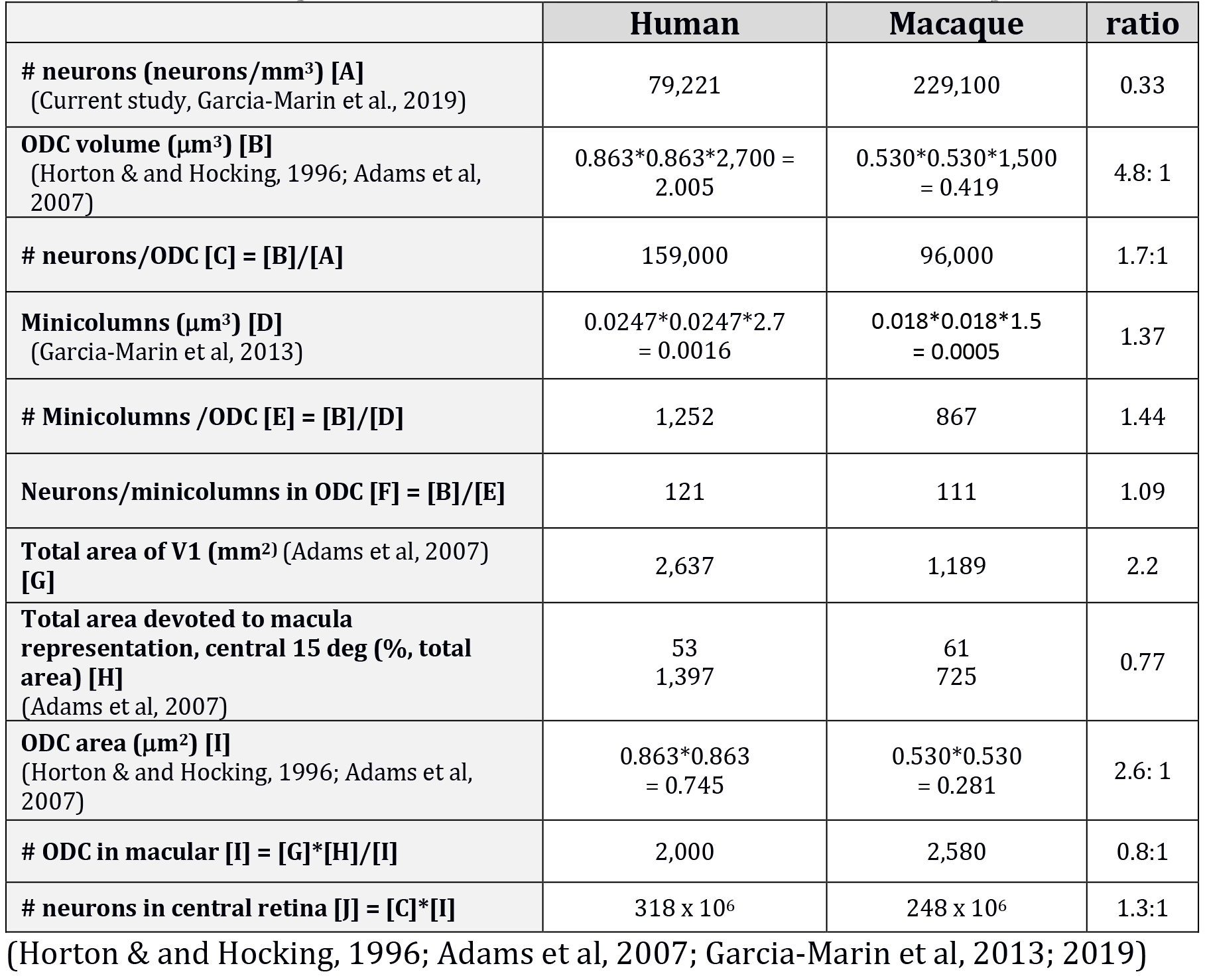
Relationship between neurons and ODC in human and macaque

|  | **Human** | **Macaque** | **ratio** |
| --- | --- | --- | --- |
| **# neurons (neurons/mm<sup>3</sup>) [A]**<br>(Current study, Garcia-Marin et al., 2019) | 79,221 | 229,100 | 0.33 |
| **ODC volume (μm<sup>3</sup>) [B]**<br>(Horton & and Hocking, 1996; Adams et al, 2007) | 0.863*0.863*2,700 =<br>2.005 | 0.530*0.530*1,500<br>= 0.419 | 4.8: 1 |
| **# neurons/ODC [C] = [B]/[A]** | 159,000 | 96,000 | 1.7:1 |
| **Minicolumns (μm<sup>3</sup>) [D]**<br>(Garcia-Marin et al, 2013) | 0.0247*0.0247*2.7<br>= 0.0016 | 0.018*0.018*1.5<br>= 0.0005 | 1.37 |
| **# Minicolumns /ODC [E] = [B]/[D]** | 1,252 | 867 | 1.44 |
| **Neurons/minicolumns in ODC [F] = [B]/[E]** | 121 | 111 | 1.09 |
| **Total area of V1 (mm<sup>2</sup>) (Adams et al, 2007) [G]** | 2,637 | 1,189 | 2.2 |
| **Total area devoted to macula representation, central 15 deg (% , total area) [H]**<br>(Adams et al, 2007) | 53<br>1,397 | 61<br>725 | 0.77 |
| **ODC area (μm<sup>2</sup>) [I]**<br>(Horton & and Hocking, 1996; Adams et al, 2007) | 0.863*0.863<br>= 0.745 | 0.530*0.530<br>= 0.281 | 2.6: 1 |
| **# ODC in macular [I] = [G]\*[H]/[I]** | 2,000 | 2,580 | 0.8:1 |
| **# neurons in central retina [J] = [C]\*[I]** | 318 x 10<sup>6</sup> | 248 x 10<sup>6</sup> | 1.3:1 |

(Horton & and Hocking, 1996; Adams et al, 2007; Garcia-Marin et al, 2013; 2019)

Next, we asked whether the increased number of neurons in the human ODC is because there are more minicolumns per ODC or more neurons in each minicolumn. Previously, we had measured the average minicolumn width in human and macaque V1, 24.7 μm and 18 μm, respectively (Garcia-Marin et al, 2013). Assuming a square array distribution of the minicolumns, we estimated that the number of minicolumns per ODC was 1252 for human and 867 for macaque (Fig. 5). Using the total number of neurons per ODC, we estimated that there were 121 neurons per minicolumn in each human ODC and 111 neurons per minicolumn in each macaque ODC. These estimates indicate that at the finest columnar scale – the minicolumn – there is a close match in the number of neurons between human and macaque. Yet at the mesoscale the human cortex has a greater number of neurons because there are more minicolumns per ODC than in the macaque. Furthermore, the increase in number is disproportionately distributed in the supra and infragranular layers.

**Figure 5:**
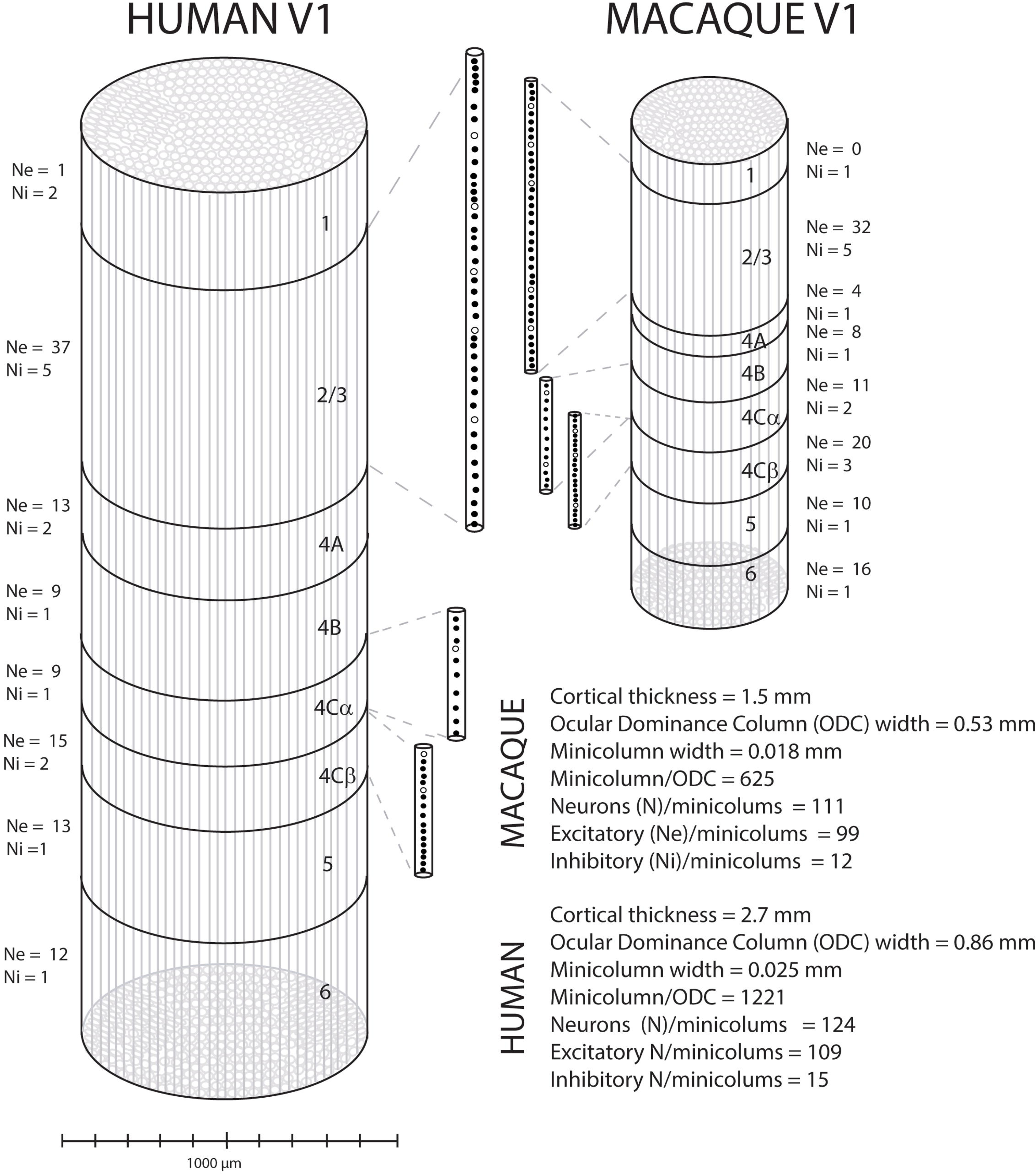
Schematic summary of the neurons within a hypercolumn (HC) in human and macaque. A hypercolumn is composed for 1221 minicolumns in human and 625 minicolumns in cynomologus macaque. For each layer the number of excitatory (Ne) and inhibitory (Ni) neurons is calculated and shown as the number of neurons in each minicolumn in each layer. Layers 2/3, 4Cα and 4Cβ are expanded twice their length and width to give a clearer impression of the distribution of neurons within a single minicolumn in each of these layers, excitatory neurons in green, inhibitory in red.

## DISCUSSION

### Areal and Laminar Neuronal Density in V1

In the current study we found that the neuronal density was about 10-15% greater than in previous studies that have used stereological methods for neuronal counting that have more than three subjects (Everall et al, 1993 – n = 13; Pakkenberg and Gundersen, 1997 – n = 63; Dorf-Petersen et al, 2007 – n = 9). There was a small difference between the male and female groups in the Pakkenberg and Gundersen (1997) study. When we compare the density measurements from males from the three studies with the results of the current study (also all males) the ratio of densities (current study/previous study) ranges from 0.84 to 0.89 (Supplementary Table 1). This confirms that the current method provides about 10-15% higher density estimates than earlier studies when taking gender differences into account. In the three earlier studies the coefficients of variation were in the range of 0.14 – 0.23, again similar to the variation found in the current study (0.13). Another study that used 3D counting regime (Selemon et al, 1995) reported substantially higher mean density (1.55 – 1.8 times higher) and lower CV (0.08) than the three studies using stereology and our current study.

One reason for this different in the density maybe the higher fidelity of specifically labeling the nuclei of neurons using the immunocytochemical labeling protocol rather than having to make a judgment about whether a Nissl stained cell is a neuron or non-neuronal cell.

There have been no previous studies that have determined laminar density where the cortex has been divided into the layer structure that includes the accepted subdivisions for layer 4 (Preuss and Colman, 2002) that match those in macaque monkey (Lund, 1973). Nor did any of the studies using stereology undertake a laminar analysis. Hence the current study is the first comprehensive description of the variation of density across layers in human V1.

There was a clear variation in density between layers (Table 2; Fig. 3A) where the highest densities were found in upper layer 2, layer 4A and layer 4Cβ (Table 3; Fig. 3B). The peak densities in these three sublayers were around 150,000 neurons/mm^3^ (Table 3), almost double the average density across all layers. In contrast, the low density regions, layer 4B and at the layer 4Cβ/5 border, had densities of just less than 40,000 neurons/ mm^3^, a density that is about half the average across cortex. These laminar variations were consistent between individuals (Table 3; Figure 3B). The mean density between individuals showed a coefficient of variation of 0.14 which is a close match to those studies using stereology (see Suppl. Table 1).

#### Glia:neuron density ratio

We found that the average non-neuronal to neuron ratio for human V1 was 1.1:1, lower than the ratio (∼1.4:1) reported with the isotropic fractionator in the cortical grey matter (Azevedo et al, 2009). Since glia, on average, represents about 70% of non-neuronal cells, we estimated that the glia to neuron ratio was about 0.76:1 in human V1. The V1 glia:neuron ratio in our current study is smaller than the values reported using stereological counting methods in the neocortex as a whole (1.3-1.7, Pelvig et al., 2008, Pakkenberg et al., 2003). It is worth noting that this glia to neuron ratio is not constant across all cortical areas; as noted by Azevedo et al (2009) the ratio changes depending on the density of neurons; the extreme example is the cerebellum where neuronal density is high and the glia to neuron ratio drops to 0.23. V1 has a neuronal density that is greater than other cortical areas therefore, following the arguments of Azevedo et al (2009), it might be expected that V1 has a lower glia to neuron ratio than the neocortex as a whole, as our results demonstrate. The glia to neuron ratio also varied through the thickness of cortex. We found an inverse relationship between neuron density and glia:neuron ratio (Suppl. Fig 1B). This largely reflects that glial and non-neuronal densities are relatively constant through the thickness of cortex (Suppl. Fig. 1A) while neuron density shows local peaks and valleys (Fig. 3B, Suppl. Fig. 1A).

The non-neuronal:neuronal ratio has also been determined in different species of macaque of V1 with a ratio of 0.44 (Giannaris and Rosene, 2012; Kelly and Hawken, 2017). This is considerably lower than the ratio we found in human in the current study in human V1 (1.1). If we assume that about 70% of the non-neuronal cells are glial cells, as we have in human, then the ratio of glia:neuron is 0.3 in macaque V1, using the same methods as in the current study (Kelly and Hawken 2019). The glia-to-neuron ratios in macaque V1 have also been reported in earlier studies, with values that range between 0.4 to 0.6 (O’Kusky and Colonnier 1982; Romme Christensen et al. 2007; Collins et al. 2010a; Giannaris and Rosene 2012). Contrary to what we observed in human, this glial:neuron ratio in macaque V1 is similar to the glia:neuron ratio in the whole cortex (0.56, Romme Christensen et al., 2007) suggesting a constant glial density across different cortical areas.

The difference between the glia:neuron ratio in human and macaque V1 (0.75 vs 0.33, respectively), when the same methods are used, indicatesthat there is an increase in the glial density in human V1. Glial cells, particularly astrocytes, play a crucial role in the flux of energy substrates to neurons by regulating the rate of glucose uptake and phosphorylation in response to glutamate concentrations in the synaptic cleft (Tsacopoulos and Magistretti, 1996).

#### Density of PV neurons across layers

Several studies have assigned a critical role to inhibitory neurons in controlling the flow of activity within local circuits, including maintaining excitatory/inhibitory balance (van Vreeswijk and Sompolinsky, 1996), shaping receptive fields (Sillito, 1975; review in Ferster and Miller, 2000), gain control (Atallah et al, 2012; Wilson et al, 2012), and surround suppression (Bair et al, 2003; Angelucci and Bressloff, 2006; Adesnik et al, 2012). For a comprehensive understanding of the interaction between excitation and inhibition in the neocortex, it is crucial to obtain data on the distribution of excitatory and inhibitory neurons in a cortical column and within a cortical layer. The fixation protocol that was used for the human postmortem tissue was incompatible with GABA immuno-histochemistry and consequently we were not able to measure the total density of GABAergic interneurons directly. PV-ir interneurons account for the largest subpopulation of the inhibitory interneuron population in rodent somatosensory cortex (Rudy et al, 2011), as well as in human and macaque V1 (Leuba et al, 1998; Kelly et al, 2019). We found that the average density of the PV population was 4.6 x 10^3^ neurons/mm^3^, a value that accounts for ∼6% of the total neuronal population. Similar percentages of PV had been reported previously the Catarrhines, including human and macaque, in different areas (Leuba et al, 1998; Gabbot and Somogyi, 1996; Glezer et al, 1998; Smiley et al, 2016; Kelly et al, 2019; Sherwood et al, 2007). If the PV population makes up approximately 50% of the total inhibitory population, as is the case for macaque and human V1 (Leuba et al, 1998; Kelly et al, 2019), then we estimate that the total proportion of inhibitory neurons would be about 12%. Recently, we estimated the density of GABAergic neurons in macaque V1 and found that they represent 11% of the total neuronal population. In human V1, Leuba et al, (1998) quantified the density of the other calcium-binding proteins, Calbindin (CB) and Calretinin (CR). Comparing their GABAergic density (from their measurements of PV, CB and CR) with our neuronal density, interneurons would represent 9.3% of the total neuronal population. Within human V1 the data indicates that the total inhibitory neuronal population is between 9 and 12% of the total neuronal density. Hence our numerical estimates provide quite narrow bounds on the density of inhibitory neurons in V1. Nonetheless, caution should be taken when extrapolating these numbers to other cortical areas, as it has been reported that the PV-ir neurons do not represent the largest inhibitory subpopulation in prefrontal or primary auditory areas in primates (Conde et al, 1994; Smiley et al, 2016; Fish et al, 2018).

#### Interareal Comparisons

In catarrhines in particular and all mammals in general the relative density in different cortical areas is thought to be of central importance from a number of views on how cortex functions. From a developmental view-point the neuronal density in each cortical area provides a strong prediction of the likely connectivity profile of that area with other cortical areas (Finlay et al, 2001). When the predications are made within this framework they are, by necessity, made within a single species of primate as the neuronal density expressed in neurons/mm^3^ of cortex shows a wide range between species. Within the Catarrhines, we have shown the average V1 neuronal density in macaque monkey is 230 x 10^3^ neurons/mm^3^ (Kelly and Hawken, 2017; Garcia-Marin et al, 2019; Table 6) and in human the density is 79 x 10^3^ neurons/mm^3^ (Table 5). Nonetheless, the regional areal variation within a species is the important factor when considering the developmental connectomic viewpoint (Findlay et al, 2001; Barbas 2015; Atapour et al, 2019). From this perspective there are currently few studies of human cortex that provide a comprehensive account of the regional areal densities of neurons. Using the methods in the current paper would provide a basis for such a series of studies.

#### Functional Considerations

A complementary view of the importance of cortical density is in terms of cortical function (Carlo and Stevens, 2013, Srinivasan et al, 2015). In the visual system of the Catarrhines there is a very clear homology across the different members of the parvorder that includes humans, non-human apes and macaque monkeys. In the cortex the commonly used measure of neuronal density (neurons/mm^3^), that works well for the within species areal comparison from the developmental view-point, is not well suited for a comparison of cortical function, especially when the comparison is between species (Herculano-Housel et al, 2008). Functionally, the columnar organization of cortex has been a fruitful viewpoint. Here we argue that one of the most frequently adopted columnar measures (Rockel et al, 1980; Dombrowski et al, 2001; Carlo & Stevens, 2013; Srinivasan et al, 2015; Atapour et al, 2019), the neuronal population under a mm^2^ column of cortex from the pial surface to the white matter, is not the most appropriate measure when making comparisons across species from a visual-processing functional viewpoint.

#### Neural Populations in an Ocular Dominance Column

As clearly discussed by Rakic (2008) there are numerous types of columns or modules. In mammalian vision a module that has been of interest since its discovery and subsequent elaboration is the ocular dominance column (ODC) (Hubel and Wiesel, 1962; 1968; Hubel et al, 1978; Horton and Hedley-Whyte, 1984; Adams et al, 2007) that is thought to contain the circuits for the emergence of orientation preference and spatial processing of a point image (Carlo and Stevens, 2013), in many mammalian species. In the current study we determined the population of neurons within each layer underlying an eye dominance module to compare with the size of the populations underlying the same module in macaque (Garcia-Marin et al, 2019) thereby providing a comparison based on functional similarity rather than evaluating the uniformity (Rockel et al, 1980; Carlo and Stevens, 2013) or nonuniformity (Herculano-Houzel et al, 2008; Lent et al, 2012) of cortex hypotheses.

The lattice of the ODCs can be determined from CO-stained sections of flatmounted V1 and the average size of the OD domain can also be determined. The area occupied by the macaque OD domain is smaller than that of the human, the ratio of areas is about 0.25:1. In cynomolgus macaque the ODC diameter is, on average, 0.53 mm (Horton and Hocking, 1996) while in humans it is considerably larger, about 0.86 mm (Adams et al, 2007). In human the number of neurons through the thickness of cortex in a single ODC is 159 x 10^3^ (Table 5) compared to 96 x 10^3^ for the similar column in macaque (Table 6). Using the ODC as a core processing unit, human cortex has 1.6 times more neurons than macaque for undertaking the same local computations (Fig. 5).

#### Neural Populations in Central Vision

The next question we addressed was the number of ODCs in a large spatial extent of visual cortex and their underlying neural populations. Often, the visual cortex is divided into the central and peripheral regions (Adams et al, 2007). In this parcellation, the central region is termed the macula region – the central 15 ° of eccentricity from the fovea to the optic disc. The remaining cortex contains a representation of the peripheral visual field, including the monocular crescent. The total area of V1 in the human is 2.2 times larger than in macaque (2,637 vs 1,189 mm^2^, respectively) (Adams et al, 2007). When comparing the macula representation of the central 15 degrees, 53% of the striate cortex is devoted to this central representation in humans versus 61% in macaques (Adams et al, 2007). As a result, in the macula region, macaques have more ODC than humans (2,580 vs 2,000 ODCs, macaque and human respectively). Each single human ODC has 1.7 times more neurons than each macaque ODC (Tables 5 and 6), however since humans have fewer ODCs in the same central region, humans only have 1.28 times more neurons in the central 15 ° than macaque (318 x 10^6^ neurons vs 248 x 10^6^ neurons, human and macaque, respectively). Therefore, the number of neurons available for the first stage of cortical processing in central vision is only ∼28% greater in humans than macaques, yet the difference in volume is 4 times greater in human than macaque. Next we address the likely sources of the additional volume in human cortex.

#### Minicolumns, Soma Size and Neuropil

We found that the number of minicolumns in each ODC was about double in human compared to macaque while the number of neurons in each minicolumn was nearly constant: 111 neurons in macaque and 120 in human. Because individual minicolumns in humans and macaques have about equal numbers of neurons and the volume of each minicolumn is 4 times larger in human than macaque (Figure 5), it is likely that the perikaryon is expanded in human cortex and the additional neuropil volume could be occupied with other elements, dendrites, synapses, and glial cells Analysis of Lucifer Yellow labeled pyramidal cells in tangential sections of layer 3 of macaque V1 and V2 demonstrated that the cross-sectional area of the somata of V1 neurons are smaller than those of V2 neurons (118 vs 130 µm^2^, respectively) (Olga et al, 2017; Elston et al, 2005; Elston and Rosa, 1998). Although there is no published data about the neural characteristics of human V1, preliminary results show that somata are smaller in V1 than in V2 (153 vs 227 µm^2^, respectively (Benavides-Piccinone et al., 2023 in preparation; Elston et al, 2001 unpublished results). If the difference in soma size of the layer 3 neurons between human and macaque V1 (1.3 times larger) holds for all of V1 cortex, it can be concluded that part of the neuropil volume in human is occupied by larger neurons than macaque neurons. The same Lucifer Yellow studies that measured the soma area also compared the basal dendritic field areas between human and macaque. These studies showed that the basal dendritic field area of the labeled layer 3 pyramidal neurons was double in human compared to macaque in V2 (Elston et al, 2001). Although there is similar data reporting the basal dendritic field area of macaque V1 neurons (Elston and Rosa, 1998, Elston et al, 2001; Oga et al, 2017), there are no reported measurements from human V1, as yet, to compare with macaque V1. However, if it is the case that the dendritic arbor is larger in human V1 than in macaque V1 as it is for V2, we can conclude that both neural soma volume and dendritic arbor size are expanded in human to occupy the larger proportion of the neuropil. More studies are needed to determine if there are also more synapses in the human neuropil than in the macaque neuropil.

To conclude, comparative anatomical studies between human and macaque are important as both species are very similar in their visual perceptual capacity (e.g De Valois et al, 1974; Kiorpes, 2016; Dahl et al, 2009; Furtak et al., 2022; Horwitz, 2015). Having an accurate estimate of the number of neurons in human V1 is a critical step to determine coding capacities of this area. When a similar visual processing region is compared between humans and an animal model it will be essential to make rigorous comparisons of the cellular distributions (as in the current study) and their functional characteristics to gain insights into their processing capacity and how they may be altered in dysfunction (Selemon et al, 1995; Dorph-Petersen et al, 2007).

---

**Supplementary Figure 1:**
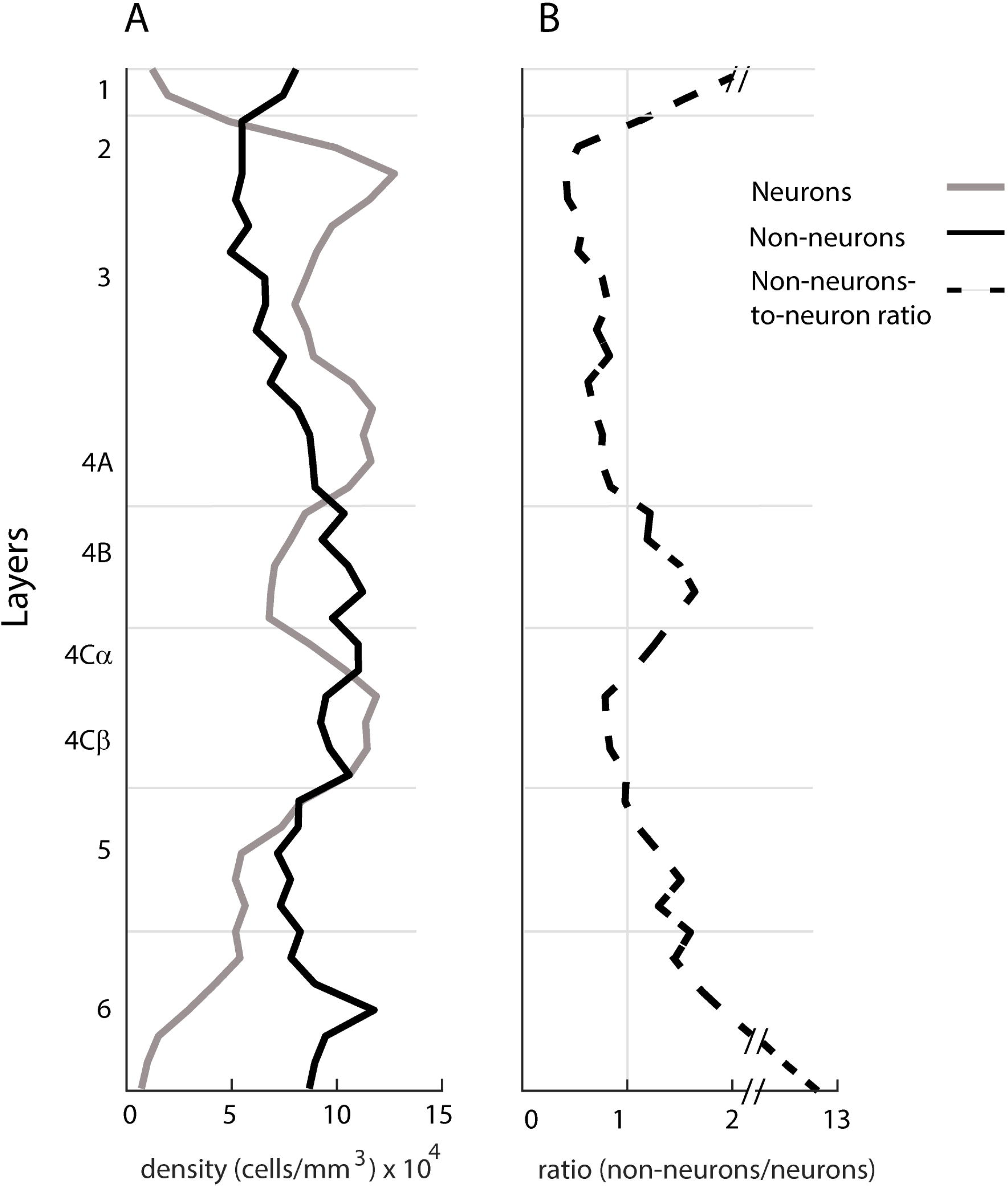
Distribution of neurons and non-neurons through the depth of human V1. (A) Continuous laminar distribution of the neurons (black) and non-neuronal cells (gray). (B) Continuous laminar profile of non-neuron to neuron ratio (dashed line).

**Supplementary Figure 2:**
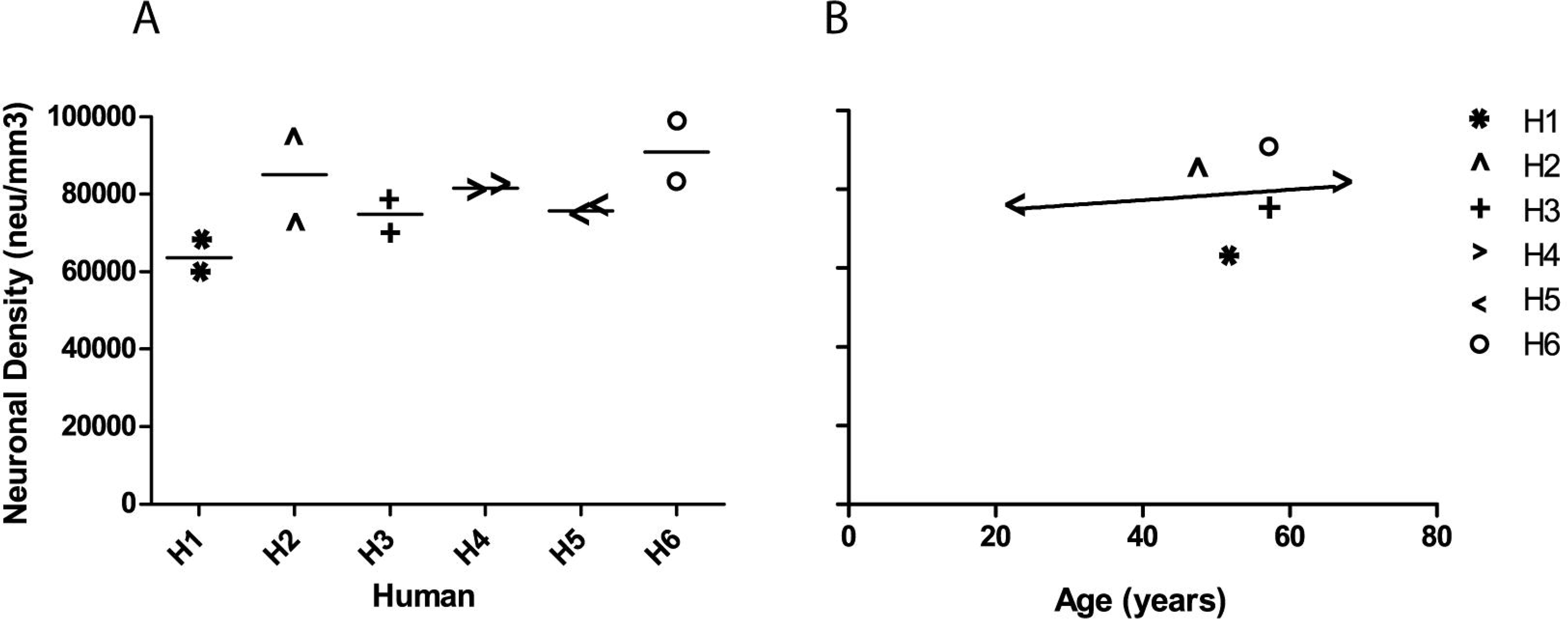
A) Average neuronal density for each human. B) Average neuronal density for each human by age.

**Supplementary Table 1:**
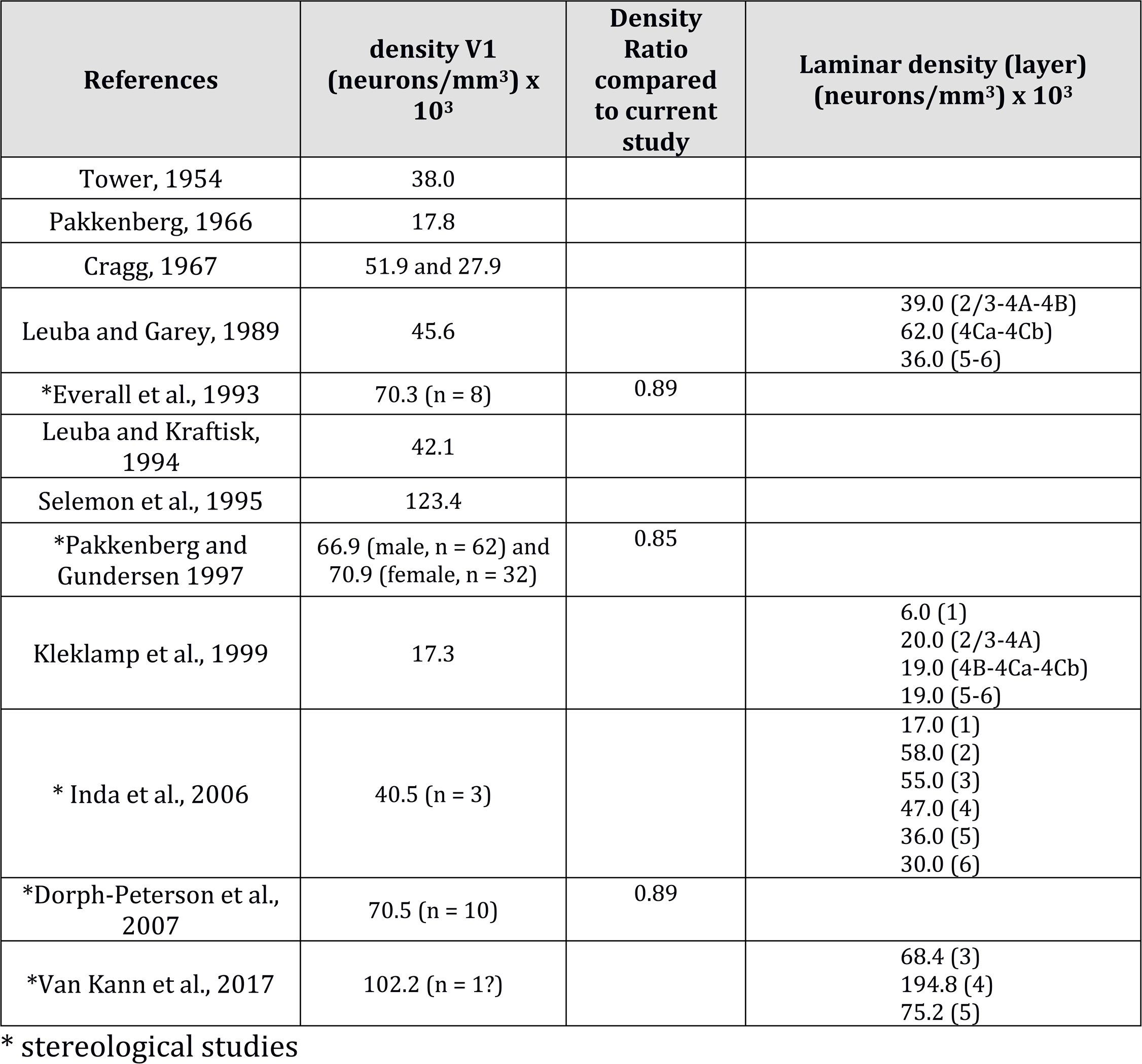
Previous data on the neuronal density in human V1 across all layers or by individual layers

| References | density V1<br>(neurons/mm<sup>3</sup>) x<br>10<sup>3</sup> | Density<br>Ratio<br>compared<br>to current<br>study | Laminar density (layer)<br>(neurons/mm<sup>3</sup>) x 10<sup>3</sup> |
| --- | --- | --- | --- |
| Tower, 1954 | 38.0 |  |  |
| Pakkenberg, 1966 | 17.8 |  |  |
| Cragg, 1967 | 51.9 and 27.9 |  |  |
| Leuba and Garey, 1989 | 45.6 |  | 39.0 (2/3-4A-4B)<br>62.0 (4Ca-4Cb)<br>36.0 (5-6) |
| *Everall et al., 1993 | 70.3 (n = 8) | 0.89 |  |
| Leuba and Kraftisk,<br>1994 | 42.1 |  |  |
| Selemon et al., 1995 | 123.4 |  |  |
| *Pakkenberg and<br>Gundersen 1997 | 66.9 (male, n = 62) and<br>70.9 (female, n = 32) | 0.85 |  |
| Kleklamp et al., 1999 | 17.3 |  | 6.0 (1)<br>20.0 (2/3-4A)<br>19.0 (4B-4Ca-4Cb)<br>19.0 (5-6) |
| * Inda et al., 2006 | 40.5 (n = 3) |  | 17.0 (1)<br>58.0 (2)<br>55.0 (3)<br>47.0 (4)<br>36.0 (5)<br>30.0 (6) |
| *Dorph-Peterson et al.,<br>2007 | 70.5 (n = 10) | 0.89 |  |
| *Van Kann et al., 2017 | 102.2 (n = 1?) |  | 68.4 (3)<br>194.8 (4)<br>75.2 (5) |

\* stereological studies

**Supplementary Table 2:** Individual case information.

| Case no. | Race | Sex | Age, y | PMI, h | COD |
| --- | --- | --- | --- | --- | --- |
| H1 | W | M | 52 | 9 | Pulmonary, thromboembolism |
| H2 | W | M | 48 | 19.8 | ASCVD |
| H3 | W | M | 57 | 22.4 | Hypertensive & ASCVD |
| H4 | W | M | 66 | 27.3 | Blunt force trauma of the trunk |
| H5 | B | M | 23 | 18.5 | Blunt force trauma of the trunk |
| H6 | B | M | 57 | 15.2 | Pulmonary, thromboembolism |
| H7 | B | M | 59 | 5.9 | cardiac tamponade |

W, white; B, black; M, male; y, year; h, hour; PMI, post mortem interval, ASCVD: atherosclerotic cardiovascular disease.

**Supplementary Table 3:** Statistical analysis of the neuronal density by layer (One-way Anova) (* < 0.05, ** < 0.01, *** < 0.001, ns, no significant)

| | L1 | L2/3 | L4a | L4b | 4Cα | 4Cβ | 5a | 5b | 6a | 6b |
| --- | --- | --- | --- | --- | --- | --- | --- | --- | --- | --- |
| L1 |  | *** | *** | ns | *** | *** | *** | * | *** | ns |
| L2/3 |  |  | ns | *** | ns | ** | ns | *** | ns | *** |
| L4a |  |  |  | *** | * | ns | ns | *** | *** | *** |
| L4b |  |  |  |  | * | *** | *** | ns | ns | ns |
| L4Cβ | | | | | | *** | ns | ns | ns | *** |
| L4Cβ | | | | | | | ** | *** | *** | *** |
| L5a |  |  |  |  |  |  |  | *** | ns | *** |
| L5b |  |  |  |  |  |  |  |  | ns | ns |
| L6a |  |  |  |  |  |  |  |  |  | ** |
| L6b |  |  |  |  |  |  |  |  |  |  |

